# No Evidence for a Difference in 2D:4D Ratio between Youth with Elevated Prenatal Androgen Exposure due to Congenital Adrenal Hyperplasia and Controls

**DOI:** 10.1101/2020.05.07.082529

**Authors:** Gideon Nave, Christina M. Koppin, Dylan Manfredi, Gareth Richards, Steven J. Watson, Mitchell E. Geffner, Jillian E. Yong, Robert Kim, Heather M. Ross, Monica Serrano-Gonzalez, Mimi S. Kim

## Abstract

The second-to-fourth digit ratio (2D:4D) has been associated with sexual dimorphism, with a lower 2D:4D in males. A large body of research has relied on the 2D:4D as a proxy for prenatal androgen exposure, and includes reports of relationships between 2D:4D and a wide range of human traits. Here, we examine the validity of the 2D:4D proxy by studying the association between 2D:4D and classical Congenital Adrenal Hyperplasia (CAH) due to 21-hydroxylase deficiency, a condition characterized by excessive prenatal exposure to androgens during most of the gestational period. To this end, we retrospectively examine 513 serial radiographs of the left hand obtained clinically in 90 youth with classical CAH (45 female) and 70 control youth (31 female). Replicating previous reports, we observe associations of the 2D:4D with sex (lower 2D:4D in males) and age (increase of 2D:4D through development). However, we find no evidence for differences in 2D:4D between CAH and controls (full sample: □ = -0.001 (−0.008, 0.006)]; females: □ = -0.004 [-0.015, 0.007]; males: □ = 0.001, [-0.008, 0.011]). Although our findings do not rule out a small association between the 2D:4D and CAH, they cast doubt on the usefulness of the 2D:4D as a biomarker for prenatal androgen exposure in behavioral research.

## Introduction

The sex hormone testosterone plays a crucial role in the prenatal sexual differentiation. A growing body of research suggests that testosterone exposure *in-utero* also affects the development of psychological and behavioral traits later in life (Bütikofer et al., 2019; Constantinescu and Hines, 2012; Knickmeyer and Baron-Cohen, 2006; Tapp et al., 2011). Studying the prenatal effects of testosterone is difficult, because measuring hormone levels *in-utero* requires high-risk procedures such as amniocentesis (Keelan et al., 2012). In an attempt to overcome this challenge, the ratio between the second (index) and fourth (ring) fingers (2D:4D) (Manning et al., 1998) has been proposed to approximate prenatal testosterone exposure. This ratio is on average lower in males compared to females (Baker, 1888; Hönekopp and Watson, 2010; Manning et al., 2004).

As of 2019, the number of published studies associating 2D:4D with later-life psychological and behavioral traits exceeds 1,400 (Leslie, 2019). The literature includes reports of associations between 2D:4D and a wide range of outcomes, including aggressive and violent behavior (Bailey and Hurd, 2005; Turanovic et al., 2017, though see Pratt et al., 2016), risk taking (Stenstrom et al., 2011, though see Neyse et al., 2020; van Leeuwen et al., 2020), empathic accuracy (Nitschke and Bartz, 2019, though see Nadler et al., 2019) and even success records of day traders (Coates et al., 2009) and sumo wrestlers (Tamiya et al., 2012).

Despite the growing literature, the validity of 2D:4D as a proxy for prenatal testosterone exposure is not well-established. Studies that directly manipulated prenatal androgen exposure in mice found conflicting evidence for causal effects of such manipulations on 2D:4D ratios (Zheng and Cohn, 2011; Huber et al., 2017).^1^ Human studies of the associations between 2D:4D and hormonal measures from amniotic fluid, umbilical cord blood, or the maternal circulation during pregnancy have found inconclusive support for the proxy’s validity (for review, see Richards, 2017). There is also evidence that 2D:4D is influenced by factors that are unrelated to prenatal androgen exposure. For example, studies of females with complete androgen insensitivity syndrome (CAIS) found that the variance in 2D:4D was similar between CAIS and controls despite complete insensitivity to androgenic influences in the former population, indicating that the lion’s share of variance in 2D:4D is not related to androgen exposure (Berenbaum et al., 2009; van Hemmen et al., 2017). Furthermore, 2D:4D has been found to increase during early-life development, demonstrating that the ratio is not prenatally fixed (McIntyre et al., 2005; Richards et al., 2017; Trivers et al., 2006).

Important evidence in support of the 2D:4D proxy’s validity comes from studies of patients with Congenital Adrenal Hyperplasia (CAH) due to 21-hydroxylase deficiency. CAH is the most common primary adrenal insufficiency in children. The resulting cortisol deficiency leads to an overproduction of adrenal androgens by the 7^th^ gestational week in classical (severe) CAH, secondary to disrupted steroid biosynthesis (Speiser et al., 2010; White and Speiser, 2002). Such significant changes in the intrauterine environment during early development could adversely impact fetal programming with organizational effects from excess androgens. This is seen in females with classical CAH, who are born with virilized, ambiguous genitalia and can exhibit male-typical play preferences in childhood (Berenbaum and Beltz, 2011; Hines, 2008; Pasterski et al., 2011). Thus, females with classical CAH are born with outward signs of androgen exposure, unlike males with classical CAH, and differential effects can be seen for behavior and fertility (Berenbaum et al., 2012; White and Speiser, 2002).

The first studies that compared 2D:4D between patients with CAH and controls appeared in the early 2000s (Brown et al., 2002; Ökten et al., 2002). Both studies were relatively small (less than 20 CAH cases), and both found that measures of 2D:4D from hand scans were lower (more masculinized) in both males and females with CAH compared to unaffected individuals. The first meta-analysis of the literature (Hönekopp and Watson, 2010), which included data from two additional studies (Buck et al., 2003; Ciumas et al., 2009), concluded that both males and females with CAH had lower 2D:4D compared to sex-matched controls. The meta-analytic effects were relatively large in size (Cohen’s *d* between 0.6 and 0.9), and observed for both hands.

Despite these advances, the set of studies conducted to-date is relatively small, and it is subject to important limitations. First, most studies have relied on 2D:4D measured from photocopies or hand scans. The two exceptions were studies conducted using hand radiographs by Ökten et al. (2002)^2^ and Buck et al. (2003), and neither of them reported a statistically significant association between CAH status and 2D:4D. This issue is of particular importance, because 2D:4D measures from hand scans and photocopies were shown to systematically differ from 2D:4D measures obtained directly from the hands (Robertson et al., 2008; Xi et al., 2014). This disparity is thought to arise from individual differences in the shapes of fat-pads at the fingertips, as well as the degree of pressure applied when the hands are placed on the glass plate of the copier or scanner (Caswell and Manning, 2009; Ribeiro et al., 2016; Wallen, 2009). As noted by Ribeiro et al. (2016), both the fat pads and the degree of pressure might vary by factors such as sex, sexual orientation and personality. As a result, 2D:4D measures from photocopies or hand scans might generate spurious associations between the 2D:4D and various traits that are unrelated to exposure to prenatal androgens.

A second limitation is that none of the studies comparing 2D:4D between patients with CAH and controls has accounted for the potentially confounding influence of skeletal development. This methodological detail is important because 2D:4D systematically increases during early-life development (McIntyre et al. 2005; Trivers et al. 2006), and patients with CAH can have advancement in skeletal maturation secondary to androgen exposure (Finkielstain et al., 2012). Therefore, failing to control for bone age might make the 2D:4D ratio appear larger (more feminine) in CAH patients, and mask lowering (masculinization) of the measure due to androgenic influence. Furthermore, some of the previously reported studies included samples where CAH patients and controls differed considerably in their average age (e.g. Rivas et al., 2014). As a consequence, differences in 2D:4D between patients with CAH and controls might reflect general differences in skeletal development rates or chronological age, rather than specific effects of prenatal androgen exposure.

The current work overcomes the limitations above, in the largest study to-date (N = 160, including 90 CAH cases) comparing the 2D:4D of patients with CAH and controls (both males and females). Our study relies on repeated 2D:4D measures from radiographs, allowing us to directly quantify finger bone structure and to overcome the confounding influences associated with 2D:4D measures from hand scans (Ribeiro et al., 2016). Our reliance on radiographs further allows us to systematically account for the potentially confounding effects of skeletal development by accurately measuring bone age and controlling for it statistically.

## Study Participants and Methods

This was a retrospective study approved by the Children’s Hospital Los Angeles (CHLA) Institutional Review Board (CHLA-15-00453, CCI-12-00020). Parents and/or participants gave their written informed consent, with assent from participants between the ages of 7 and 14 years. This was in compliance with the Code of Ethical Principles for Medical Research Involving Human Subjects of the World Medical Association (Declaration of Helsinki).

### Participants

Ninety youth (45 females [49.5%]; 1.1-18.7 years old at the time of imaging) with classical CAH due to 21-hydroxylase deficiency and seventy control youth (31 females [44.3%], 2.6-19.7 years old at the time of imaging) were retrospectively studied (see sample characteristics in **Table 1**). Youth with CAH were patients at the CHLA pediatric endocrinology clinic (CAH Comprehensive Care Center) who were diagnosed with CAH due to 21-hydroxylase deficiency biochemically (elevated basal and/or ACTH-stimulated serum 17-hydroxyprogesterone and androgens) and/or by *CYP21A2* genotype variant. The CAH group consisted of 76% subjects with the salt-wasting (SW) form (*N* = 69, 28 females) and 24% with the simple-virilizing (SV) form (*N* = 22, 17 females) based on clinical phenotype and/or genotype.

**Table 1:** Sample and Image characteristics (standard deviations in parenthesis)

| Sample and Image Characteristic | CAH | Control |
| --- | --- | --- |
| Number of Individuals | 90 | 70 |
| Number of Females | 45 | 31 |
| Hispanic/ Latino | 53 | 33 |
| Not Hispanic/ Latino |  |  |
| - Asian | 4 | 2 |
| - African | 1 | 0 |
| - White | 30 | 35 |
| - Mixed, Asian/White | 2 | 0 |
| **Image Characteristic** | **CAH** | **Control** |
| Number of Images | 413 | 119 |
| Images per individual | 4.59<br>(1.95) | 1.7<br>(1.36) |
| Chronological Age [years] | 8.57<br>(3.89) | 11.43<br>(3.70) |
| Bone Age [years] | 9.53<br>(4.14) | 10.91<br>(3.64) |
| Height [cm] | 129.48<br>(21.91) | 138.98<br>(22.57) |
| Height Normalized | -0.21<br>(1.29) | -1.22<br>(1.26) |

The control group compromised a sample of healthy participants who had been seen at the CHLA pediatric endocrinology clinic for an assessment of short stature (*N* = 34), as well as control participants from two different research studies (*N* = 13, Kim et al., 2015; and *N* = 23, Herting et al., 2020). The characteristics of these two control samples are summarized in **Supplementary Table 1**. The clinic controls were mostly non-Hispanic, and they were shorter than average. The research study controls were mostly Hispanic, older than the clinical controls and typical in terms of age-normalized height. Control participants were otherwise healthy, and they were eligible if they had benign short stature due to familial short stature and/or constitutional delay of growth. We excluded control participants if (i) they had a pathological etiology for short stature, including a genetic or chromosomal abnormality, and /or (ii) they were medically eligible for, or treated with, growth hormone.

### Measures

#### Hand Radiographs

Serial radiographs of the left hand and wrist were obtained for bone age assessment as part of standard care in youth with CAH, and as part of an evaluation of short stature in otherwise healthy control youth. A smaller proportion of CAH and control participants had x-rays obtained as part of different research studies (Kim et al., 2015; Herting et al., 2020). Our reliance on images of the left hand is a result of largely utilizing x-rays taken for clinical purposes that are typically of the left hand (similar to Buck et al., 2003). While a previous meta-analysis reported greater sex differences in 2D:4D ratios in the right hand (Hönekopp and Watson, 2010), this analysis observed such sex differences also in the left hand, as well as a fairly large effect of CAH status on left-hand 2D:4D (*d*=.75). A pediatric endocrinologist (either M.S.K. or M.S-G.) read the radiographs in a blinded fashion to calculate bone age using the Greulich-Pyle method (Gilsanz and Ratib, 2005; Greulich and Pyle, 1953). All participants in the CAH group and 32 participants in the control group (45.7%) had multiple hand radiographs taken at different ages. The total number of CAH hand radiographs was 413 (4.59 images per person; SD = 1.95, min = 1 max = 10). The total number of control hand radiographs was 119 (1.7 images per person; SD = 1.36, min = 1 max = 7). Image characteristics are summarized on **Table 1**.

#### Digit Ratios

Two independent raters measured the second and fourth finger lengths from radiographs using digital software (Rasband, 2011; Schneider et al., 2012). First, they drew horizontal lines at the metaphysis of each finger segment (proximal, middle, and distal), and located the midpoint per line. They then measured finger length by drawing a vertical line from the distal tip through the proximal epiphysis through the midpoints (Figure 1). Finally, they calculated the 2D:4D ratio by dividing the average (of the two raters) length measure of the second finger by that of the fourth finger (inter-class correlation between raters was .97, **Supplemental Figure 1**). The means and standard deviations of the 2D:4D ratio in our sample (by CAH status, sex and age groups) are summarized in **Table 2**. These values are highly similar to those observed in a previous longitudinal study that measured 2D:4D from radiographs in typically developing populations (McIntyre et al., 2005).

**Table 2:** 2D:4D summary statistics by age, sex and CAH status.

|  |  | CAH<br>Males | Control<br>Males | CAH<br>Females | Control<br>Females |
| --- | --- | --- | --- | --- | --- |
| **Ages 0-5** | Mean | 0.892 | 0.909 | 0.907 | 0.910 |
|  | SD | 0.031 | 0.015 | 0.027 | N/A |
|  | N | 36 | 8 | 43 | 1 |
| **Ages 5-10** | Mean | 0.913 | 0.915 | 0.917 | 0.927 |
|  | SD | 0.023 | 0.015 | 0.023 | 0.014 |
|  | N | 87 | 17 | 93 | 12 |
| **Ages 10-15** | Mean | 0.919 | 0.916 | 0.922 | 0.936 |
|  | SD | 0.022 | 0.019 | 0.027 | 0.023 |
|  | N | 86 | 32 | 48 | 30 |
| **Ages 15-20** | Mean | 0.926 | 0.905 | 0.920 | 0.918 |
|  | SD | 0.02 | 0.016 | 0.04 | 0.024 |
|  | N | 15 | 10 | 5 | 9 |

**Figure 1.**
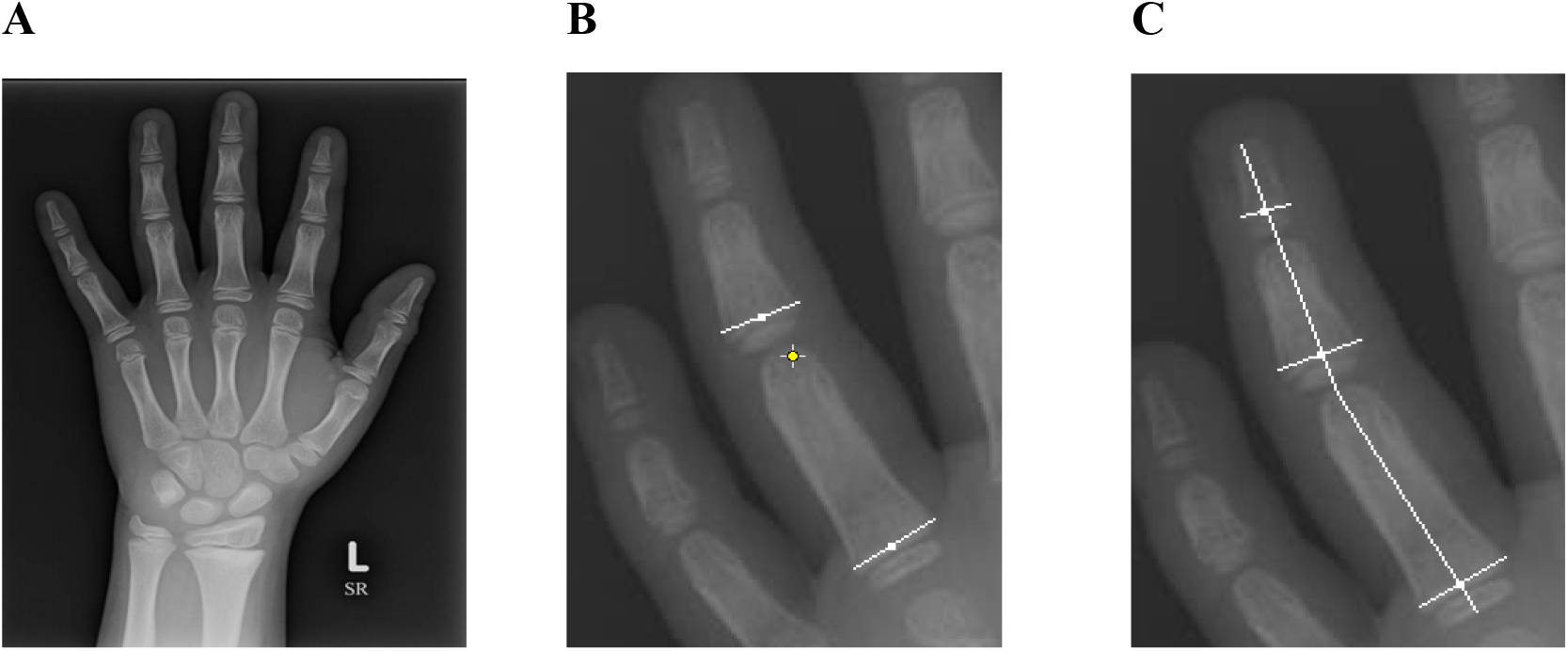
Measurements of the 2^nd^ and 4^th^ finger lengths. **(A)** Radiograph of the left hand. **(B**) Horizontal lines were drawn at the metaphysis of each phalangeal segment and the midpoint located per line (proximal, middle, and distal metaphysis). (**C**) A vertical line was drawn from the distal tip through the proximal epiphysis to give the finger length.

#### Additional Measures

Our analyses include controls for various factors that may be associated with the 2D:4D ratio based on previous findings (summarized in **Table 1**). Our primary statistical models include controls for sex (Female=1, Male=0) and bone age (in years). We also conducted secondary analyses that included additional controls for chronological age (years), height (measured with a stadiometer; cm), height normalized to chronological age and sex per United States population data^3^ (z-score), puberty status (Tanner puberty stage 1-5, assessed by a pediatric endocrinologist) and ethnicity (self-reported by participants from two prospective research studies and classified by the investigators for patients from the clinic). The most noticeable difference between the CAH and control patients were in age and normalized height, where controls were older and shorter on average (the latter difference was driven by the characteristics of the clinical controls; see **Supplemental Table 1**). The CAH group was also shorter than average relative to the general population (as indicated by a negative average normalized height), which was expected and can be attributed to androgen excess, advanced bone age, and glucocorticoid dose in CAH (Finkielstain et al., 2012). Importantly, our analysis systematically controls for bone-age, chronological age, height, normalized height and pubertal stage, in order to rule out the possible confounding influence of these variables (see further below).

The correlations between the different variables used in the study are summarized in **Table 3**. As expected, there were strong positive correlations between chronological age, bone age, height, and puberty status. This collinearity implies that while it is possible to control for these factors in our analyses, disentangling their individual contributions is not straightforward. Although bone age and chronological age were highly correlated overall, we did observe advanced bone ages in the CAH group (48.4% of patients had at least one radiograph with bone age 2 SD above the age-adjusted mean; see **Figure 2**), a finding which is in line with previous reports (Finkielstain et al., 2012).

**Table 3:**
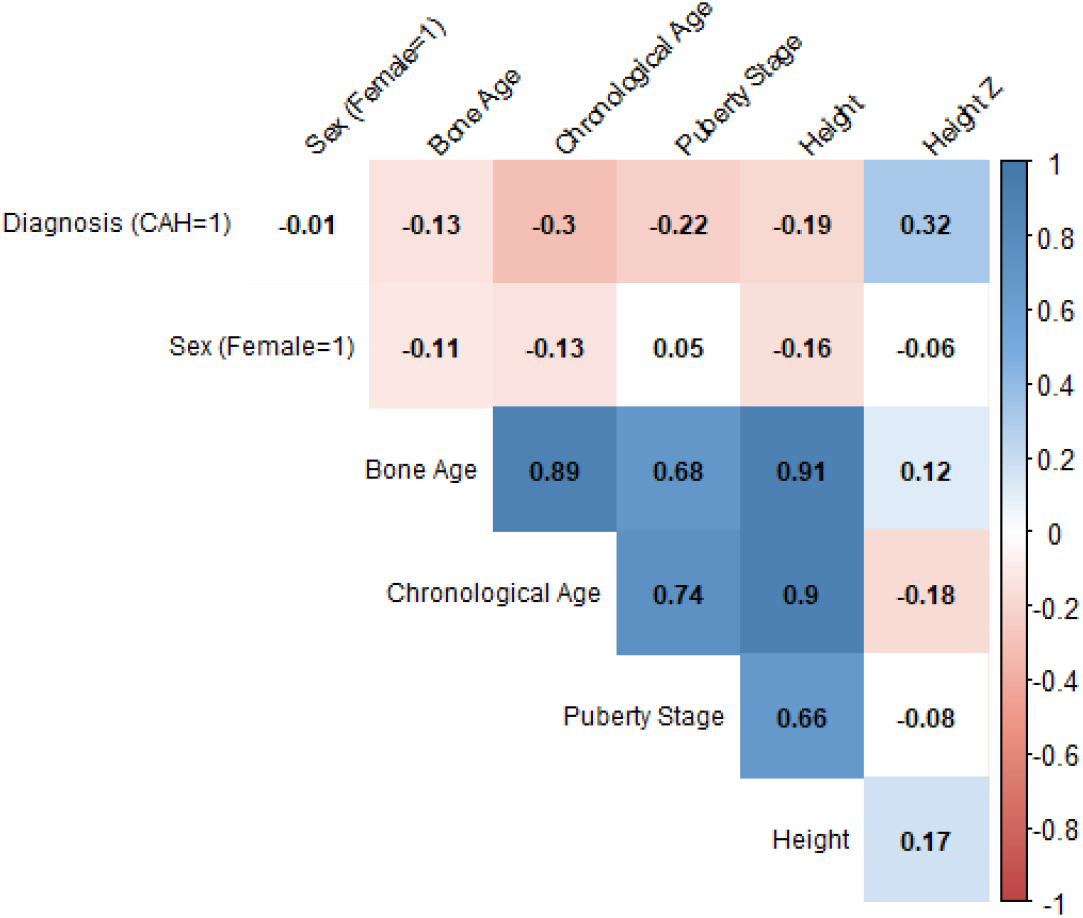
2D:4D Correlations between the main study variables.

**Figure 2.**
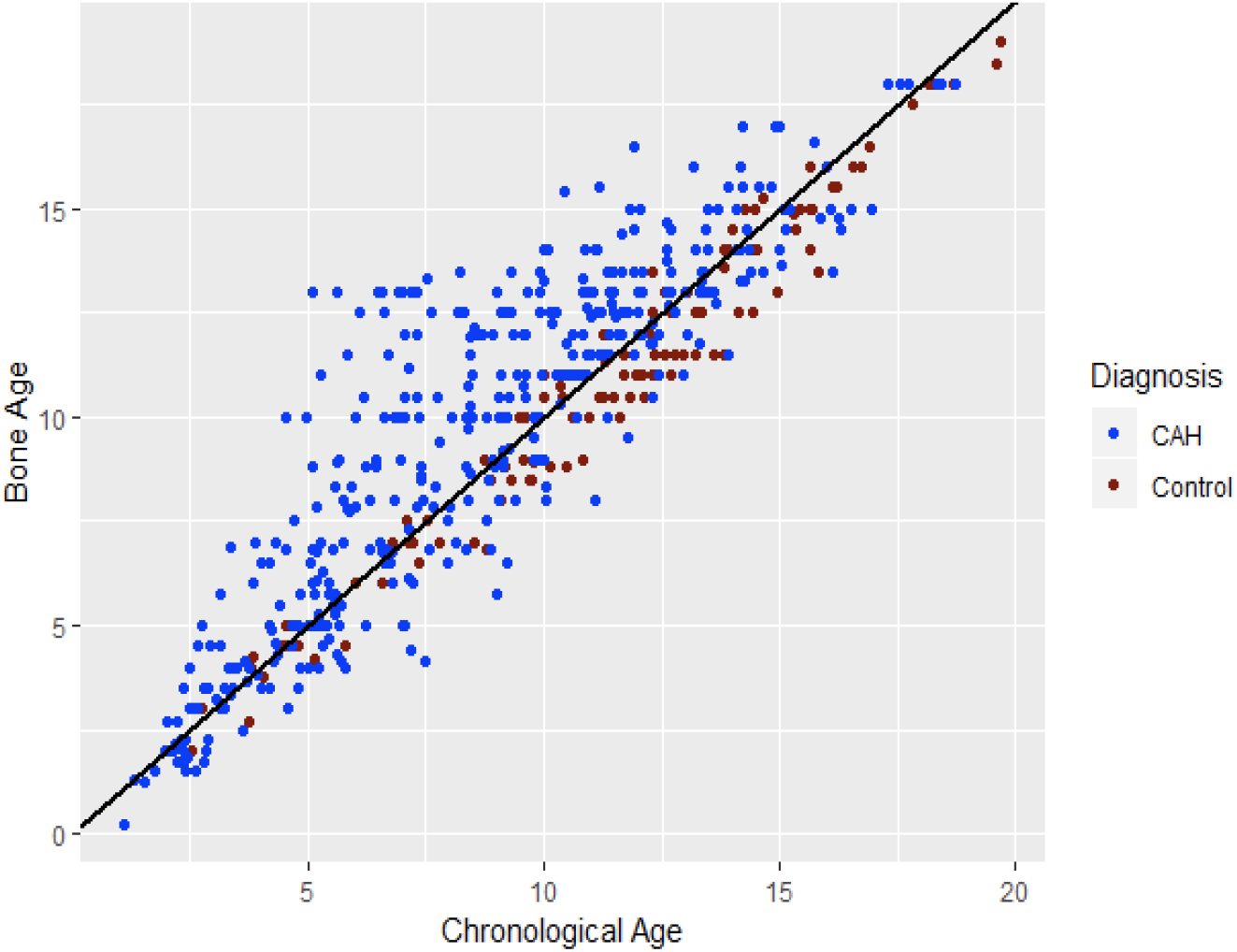
Relationship between chronological age and bone age in the study participants. The black line is a fitted regression line for the full sample.

### Statistical analyses

We estimated mixed-effects linear regression models, implemented using the ‘lme4’ package (Bates et al., 2015) in the statistical software ‘R’ to quantify the associations between 2D:4D and the explanatory variables. The outcome variable in all models was the mean left-hand 2D:4D across the two raters. Our main analysis (Model A) included fixed effects for CAH status (CAH=1; Control=0), sex and bone-age, as well as participant-specific random intercepts. We estimated additional mixed effect regressions (Models B-F) to evaluate the sensitivity of our results to different model specifications. Model B included an interaction effect between CAH status and sex, but was otherwise identical to Model A. Model C included additional controls for chronological age, height, age-normalized height, puberty status (5 categories; dummy coded) and ethnicity (3 categories: White, Hispanic/Latino, Other; dummy coded), and was otherwise identical to Model A (observations with missing values of any of the additional control variables were omitted). Models D and F were identical to Model A, but were estimated separately for females and males, respectively. Models E and G were identical to Model C, but were estimated separately for females and males, respectively.

## Results

**Table 4** summarizes the coefficient estimates of our regression models. We replicated previous findings (Hönekopp and Watson, 2010; Manning et al., 2007) that 2D:4D was greater in females (□ = 0.009, 95% confidence interval (CI) = [(0.002, 0.016)]). We also replicated the finding of McIntyre et al. (2006) that 2D:4D increases with bone age (□ = 0.002, 95% CI = [(0.001, 0.002)], **Supplemental Figure 2A**). Additionally, 2D:4D had a positive relationship with height (see **Supplemental Figure 2B)** and chronological age, though as these two variables correlated strongly with one another and with bone age, it was not possible to disentangle the differential effects of these three factors.

**Table 4:** Effects of CAH status and controls on left hand 2D:4D. Linear Mixed models with participant random intercepts. Values denote beta coefficients; 95% Confidence intervals in parenthesis

|  | Outcome variable: Left 2D:4D |  |  |  |  |  |  |
| --- | --- | --- | --- | --- | --- | --- | --- |
|  | Both<br>(Model A) | Both<br>(Model B) | Both<br>(Model C) | Females<br>(Model D) | Females<br>(Model E) | Males<br>(Model F) | Males<br>(Model G) |
| Diagnosis<br>(CAH=1) | -0.001<br>(-0.008, 0.006) | 0.001<br>(-0.009, 0.011) | -0.001<br>(-0.008, 0.007) | -0.004<br>(-0.015, 0.007) | -0.006<br>(-0.017, 0.006) | 0.001<br>(-0.008, 0.011) | 0.002<br>(-0.008, 0.012) |
| Sex (F=1) | 0.009*<br>(0.002, 0.016) | 0.012*<br>(0.0004, 0.023) | 0.009*<br>(0.002, 0.016) |  |  |  |  |
| Bone Age (years) | 0.002***<br>(0.001, 0.002) | 0.002***<br>(0.001, 0.002) | 0.002**<br>(0.001, 0.003) | 0.001***<br>(0.001, 0.002) | 0.002<br>(-0.00001, 0.003) | 0.002***<br>(0.001, 0.002) | 0.002*<br>(0.0003, 0.004) |
| Diagnosis x Sex |  | -0.005<br>(-0.019, 0.010) |  |  |  |  |  |
| Chron. Age<br>(years) |  |  | -0.004***<br>(-0.007, -0.002) |  | -0.002<br>(-0.004, 0.0004) |  | -0.011***<br>(-0.015, -0.007) |
| Height (cm) |  |  | 0.001***<br>(0.0005, 0.001) |  | 0.0003<br>(-0.00004, 0.001) |  | 0.002***<br>(0.001, 0.003) |
| Height (z) |  |  | -0.006***<br>(-0.009, -0.003) |  | -0.002<br>(-0.005, 0.001) |  | -0.015***<br>(-0.019, -0.010) |
| Puberty Stage | No | No | Yes | No | Yes | No | Yes |
| Ethnicity | No | No | Yes | No | Yes | No | Yes |
| Intercept | 0.896***<br>(0.889, 0.904) | 0.895***<br>(0.887, 0.903) | 0.834***<br>(0.808, 0.860) | 0.910***<br>(0.900, 0.920) | 0.892***<br>(0.861, 0.923) | 0.893***<br>(0.884, 0.902) | 0.738***<br>(0.691, 0.786) |
| Observations | 532 | 532 | 465 | 241 | 206 | 291 | 259 |
| Log Likelihood | 1,410.512 | 1,406.730 | 1,207.574 | 653.233 | 538.482 | 753.651 | 643.061 |
| Akaike Inf. Crit. | -2,809.025 | -2,799.459 | -2,385.149 | -1,296.466 | -1,048.964 | -1,497.302 | -1,258.122 |
| Bayesian Inf. Crit. | -2,783.365 | -2,769.523 | -2,323.018 | -1,279.042 | -1,002.374 | -1,478.935 | -1,208.326 |

Note:
\*p<.05 \*\*p<.01 \*\*\*p<.001

The regression coefficient for CAH diagnosis was not statistically distinguishable from zero under any of the model specifications in the full sample (□ = -0.001, 95% CI = [(−0.008, 0.006)]; Model A), where its point estimate was less than 10% of the effect of sex. The interaction coefficient between sex and CAH diagnosis was also not statistically distinguishable from zero (□ = -0.005, 95% CI = [-0.019, 0.010]; Model B). Analyses stratified by sex could not identify reliable effects of CAH status in either females (□ =-0.004, 95% CI = [-0.015, 0.007]; Model D) or males (□ = 0.001, 95% CI = [(−0.008, 0.011)]; Model F). Further analyses with additional controls for chronological age, height (absolute and age normalized), pubertal status and self-reported ethnicity yielded similar results (Models C, E and G). Finally, ordinary least squares (OLS) regressions that were estimated using only the last radiograph taken from each participant (and were otherwise identical to the main analysis) could not identify a significant association between CAH diagnosis and 2D:4D (see **Supplemental Table 2**).

## Discussion

We conducted a retrospective study to quantify the relationship between the 2D:4D ratio and various factors, including diagnosis of CAH, which provides a natural human model of excess prenatal androgens from early in fetal development. Our investigation, which is the largest single study of the topic to-date, relied on repeated measures of 2D:4D from finger lengths in hand radiographs. Our results replicate previously reported associations of 2D:4D with regard to sex and age. However, the associations between 2D:4D and CAH status are not distinguishable from zero, within and across sex.

Our findings are inconsistent with the results of a meta-analysis by Hönekopp & Watson (2010), which reported relatively large associations between CAH status and 2D:4D (Cohen’s *d* between 0.6 and 0.9). Of note, our study had statistical power of over 90% to detect even the smallest effect reported in the meta-analysis (d = 0.6).^4^ The discrepancy between the results of our study and the meta-analysis may arise from the fact that the latter mostly relied on studies that measured 2D:4D on outlined soft tissue from hand scans and photocopies, rather than bone measures on radiographs. Crucially, the hypothesis that initially motivated the use of the 2D:4D biomarker was that androgens directly influence skeletal development (McIntyre et al., 2007, Zheng et al. 2011), whereas measures from hand scans are also influenced by other factors that are not prenatally fixed, such as fat pads and the amount of pressure applied on the scanner surface (Berenbaum et al., 2009; Ribeiro et al., 2016; Trivers et al. 2020).

Other potential causes for this discrepancy could be differences in sample characteristics, the existence of publication bias in the meta-analyzed literature (Carter and McCullough, 2014; Lane et al., 2016) or the presence of a statistical artifact due to the small number of meta-analyzed studies (Borenstein et al., 2009). Nonetheless, our results are in line with a previous study that used 2D:4D measures from radiographs (Buck et al., 2003) that did not observe any differences between females with and without CAH. Our studies shared similar approaches, including a similar control group with shorter stature, and measurements of bone tissue from left-hand radiographs.

Our findings do not rule out the possibility of a small association between prenatal androgen elevation due to CAH and 2D:4D. This is especially true for females, for whom the point estimate of the effect of CAH on 2D:4D was negative. Nonetheless, our findings raise questions concerning the validity of behavioral research that has relied on 2D:4D to approximate prenatal testosterone exposure. This is because the relationship between 2D:4D and other phenotypes that are commonly studied in this literature is expected to be even weaker than the association that we observed between 2D:4D and CAH diagnosis. Indeed, our study examined 2D:4D in relation to a phenotype that is known to have a strong relationship with prenatal androgen exposure, particularly in females, and its phenotypic assessments were based on clinical diagnoses with virtually no measurement error. Conversely, most 2D:4D studies of typically developing populations report correlations between digit ratios and traits that are complex (e.g., aggressive behavior, risk taking, cognitive empathy). Such traits are measured with less reliability than CAH diagnosis, and are likely associated with many genetic and environmental factors that are not directly related to prenatal testosterone exposure (e.g., Aydogan et al, 2019). Furthermore, we measured 2D:4D from bone tissue on radiographs, whereas most previous studies in the literature have relied on 2D:4D measured from photocopies or hand scans. The latter are noisier measures of digit ratios, due to individual differences in finger fat tissue and the degree of pressure applied when the hands are scanned (Ribeiro et al., 2016).^5^ Finally, the much larger associations of age and height with 2D:4D that we observed suggests that factors unrelated to prenatal androgen exposure influence the 2D:4D measure, potentially confounding associations between this proxy and behavioral outcomes.

One limitation of our study is that we only used radiographs of the left hand. We therefore cannot rule out the possibility of an effect of CAH on digit ratios of the right hand. Nonetheless, it is important to note that while a meta-analysis of studies using hand scans documented greater sex differences in the 2D:4D of the right hand (*d*=.457; Hönekopp and Watson, 2010), this analysis observed sex differences also in the left hand (*d*=.376), as well as a large association between CAH status and left-hand 2D:4D. Our study, too, was able to detect significant sex differences in the left hand, yet failed to observe such differences between CAH patients and controls. Relatedly, a large study of over 3,000 radiographs found that the 2D:4D is symmetrical, with only minor differences between the hands (Robertson et al., 2008), and similarly negligible differences between the left- and right-hand 2D:4D were observed in the BBC Internet Study (the largest ever study of digit ratio; Manning et al., 2007). Given these findings, it is unlikely that any potential association between CAH status and the right hand 2D:4D would be substantially larger than the associations we observe for the left hand.

A second limitation is that our results were based on a sample of youth whose radiographs were taken for clinical purposes, rather than a random sample of the population (similar to Buck et al., 2003). Yet, our analyses systematically controlled for bone age, chronological age, height (absolute and age-normalized), ethnicity, and pubertal status. We have no reason to believe that our results depend on other characteristics of the study participants, materials, and/or context.

In summary, we do not find evidence for a relationship between 2D:4D ratio and CAH status in our sample, yet we do find that the measure is associated with other factors. This finding suggests that previously reported correlations between 2D:4D and a wide range of behavioral and cognitive traits should be interpreted with caution. We believe, however, that continued investigations of the relationships between prenatal testosterone exposure and behavioral outcomes would be worthwhile. One possibility for conducting such research is to directly study developmentally unique populations such as CAH or CAIS. Although obtaining large samples of participants from these populations might be difficult, the validity of this approach is based on a more solid empirical ground than relying on the 2D:4D ratio as a proxy for prenatal androgen exposure.

## Acknowledgments

We gratefully thank the patients and families who participated in the Natural History Study of CAH from Infancy, and the Testosterone and CAH Study at our center. We thank CARES Foundation for its support of the CHLA CAH Comprehensive Care Center (to M.S.K and M.E.G.), and the Abell Foundation for ongoing support of CAH research at our center (to M.E.G.). The writing of this paper was in part funded by K23HD084735-01A1 (to M.S.K.). The contents of this work are solely the responsibility of the authors and do not necessarily represent the official views of the National Institutes of Health. G.N. thanks the Wharton Neuroscience Initiative and Carlos and Rosa de la Cruz for ongoing support.

---

**Supplemental Figure 1.**
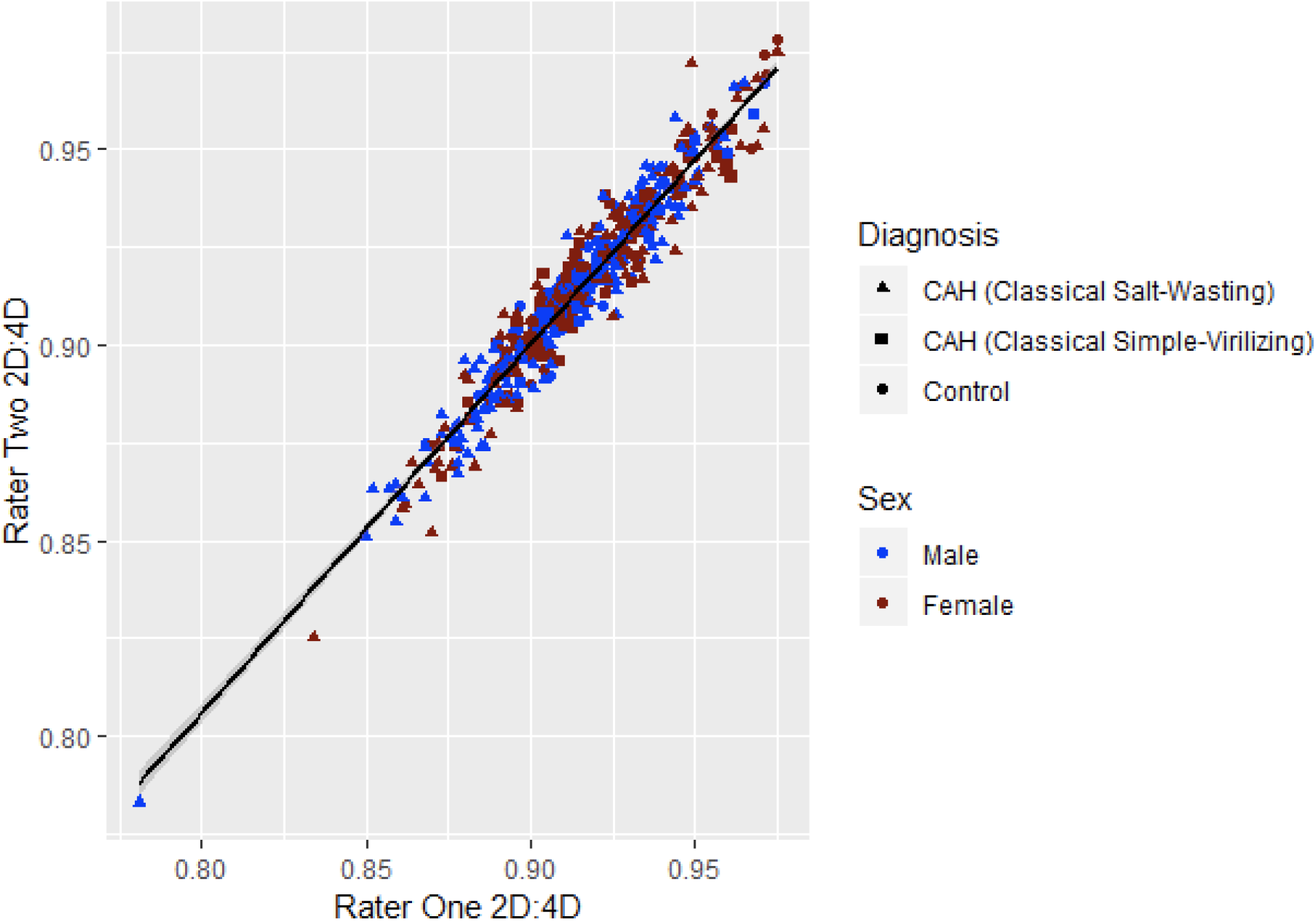
2D:4D ratio measurements in youth with and without classical CAH due to 21-hydroxylase deficiency. 413 hand radiographs in 91 youth with classical CAH (salt-wasting and simple-virilizing forms) and 119 hand radiographs in 70 controls were measured by two independent raters for 2D:4D.

**Supplemental Figure 2.**
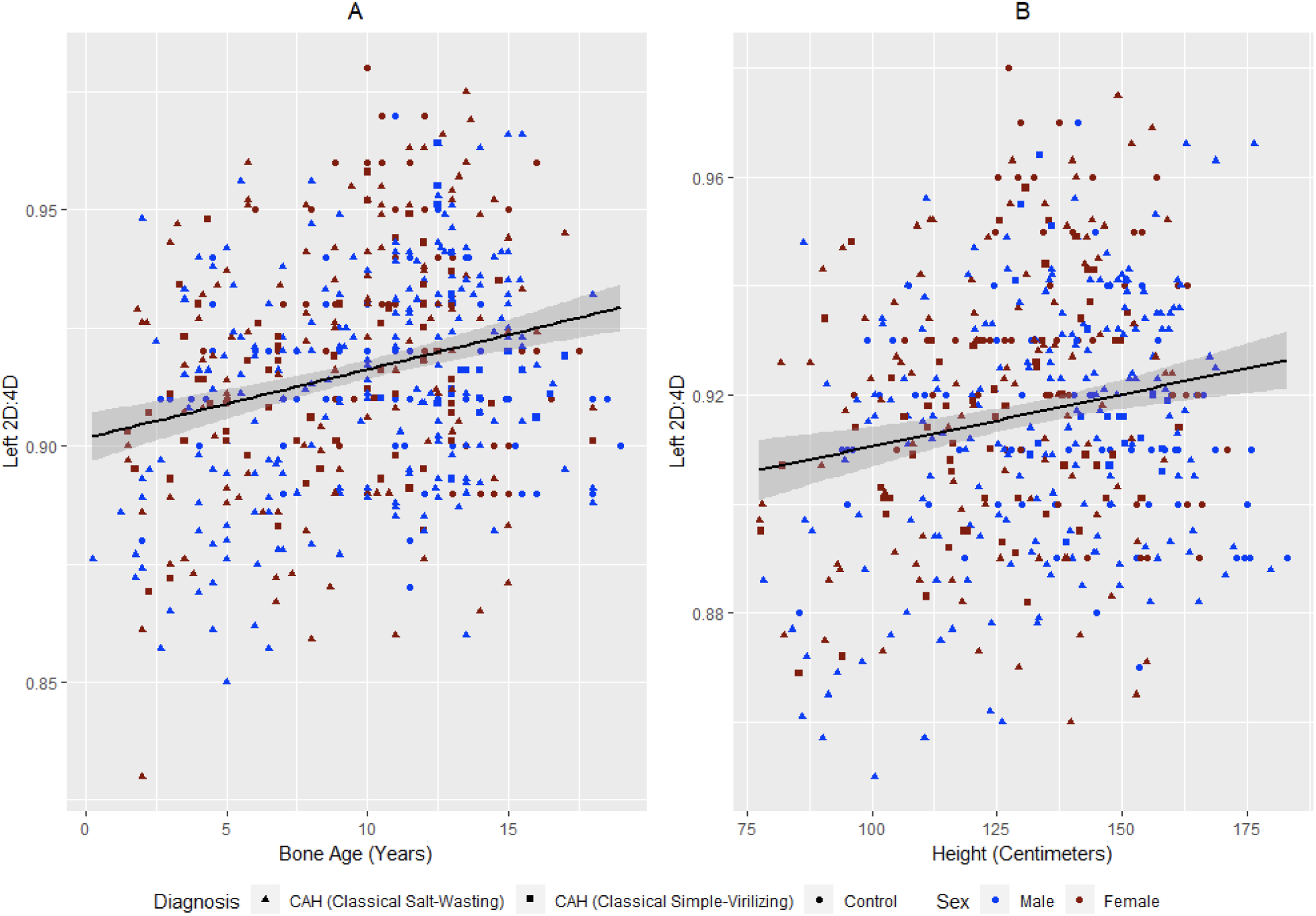
**(A)** Relationship between bone age (years) and 2D:4D of the left hand (in all participants). **(B)** Relationship between height (cm) and 2D:4D of the left hand (in all participants). The black lines are fitted regression lines.

**Supplemental Table 1:** Sample and Image characteristics (standard deviations in parenthesis), clinic and research study controls.

| **Sample and Image Characteristic** | **Clinic controls** | **Research controls** |
| --- | --- | --- |
| Number of Individuals | 34 | 36 |
| Number of Females | 12 | 19 |
| Hispanic/ Latino | 2 | 31 |
| Not Hispanic/ Latino |  |  |
| - Asian | 2 | 0 |
| - African | 0 | 0 |
| - White | 30 | 5 |
| - Mixed, Asian/White | 0 | 0 |
| **Image Characteristic** | **Clinical controls** | **Research controls** |
| Number of Images | 83 | 36 |
| Chronological Age [years] | 10.38<br>(3.37) | 13.85<br>(3.30) |
| Bone Age [years] | 9.68<br>(3.17) | 13.75<br>(3.02) |
| Height [cm] | 130.80<br>(21.24) | 156.25<br>(14.03) |
| Height Normalized | -1.80<br>(0.93) | 0.03<br>(0.95) |

**Supplemental Table 2:** Effects of CAH status and controls on left hand 2D:4D (last radiograph of each participant), OLS Regressions; Values denote beta coefficients; 95% Confidence intervals in parenthesis.

| *Outcome variable: Left 2D:4D* |  |  |  |  |  |  |  |
| --- | --- | --- | --- | --- | --- | --- | --- |
|  | (Model A) | (Model B) | (Model C) | (Model D) | (Model E) | (Model F) | (Model G) |
|  | Both | Both | Both | Females | Females | Males | Males |
| **Diagnosis (CAH=1)** | -0.0005<br>(-0.008, 0.007) | 0.004<br>(-0.005, 0.014) | -0.003<br>(-0.011, 0.006) | -0.006<br>(-0.017, 0.005) | -0.010<br>(-0.023, 0.004) | 0.004<br>(-0.005, 0.014) | 0.006<br>(-0.006, 0.018) |
| **Sex (F=1)** | 0.006<br>(-0.001, 0.013) | 0.011*<br>(0.001, 0.022) | 0.006<br>(-0.002, 0.014) |  |  |  |  |
| **Bone Age (years)** | -0.001<br>(-0.002, 0.0002) | -0.001<br>(-0.002, 0.0001) | 0.0004<br>(-0.003, 0.003) | -0.001<br>(-0.002, 0.001) | 0.0003<br>(-0.005, 0.005) | -0.001<br>(-0.002, 0.0004) | 0.0004<br>(-0.004, 0.004) |
| **Diagnosis x Sex** |  | -0.010<br>(-0.025, 0.004) |  |  |  |  |  |
| **Chron. Age (years)** |  |  | -0.002<br>(-0.005, 0.002) |  | -0.002<br>(-0.007, 0.003) |  | -0.004<br>(-0.012, 0.004) |
| **Height (cm)** |  |  | 0.0002<br>(-0.0002, 0.001) |  | 0.0001<br>(-0.0003, 0.001) |  | 0.001<br>(-0.001, 0.002) |
| **Height (z)** |  |  | -0.003<br>(-0.007, 0.001) |  | -0.002<br>(-0.007, 0.003) |  | -0.008<br>(-0.020, 0.003) |
| **Diagnosis (CAH=1)** |  |  |  |  |  |  |  |
| **Puberty Stage** | No | No | Yes | No | Yes | No | Yes |
| **Ethnicity** | No | No | Yes | No | Yes | No | Yes |
| **Intercept** | 0.926***<br>(0.912, 0.940) | 0.925***<br>(0.911, 0.939) | 0.910***<br>(0.866, 0.953) | 0.936***<br>(0.916, 0.956) | 0.925***<br>(0.870, 0.980) | 0.925***<br>(0.908, 0.943) | 0.855***<br>(0.726, 0.985) |
| **Observations** | 160 | 160 | 154 | 76 | 71 | 84 | 83 |
| **R<sup>2</sup>** | 0.034 | 0.047 | 0.101 | 0.029 | 0.204 | 0.027 | 0.130 |
| **Adjusted R<sup>2</sup>** | 0.016 | 0.022 | 0.024 | 0.002 | 0.056 | 0.003 | -0.005 |
| **Residual Std. Error** | 0.023 (df = 156) | 0.023 (df = 155) | 0.023 (df = 141) | 0.023 (df = 73) | 0.023 (df = 59) | 0.022 (df = 81) | 0.022 (df = 71) |
| **F Statistic** | 1.850 (df = 3; 156) | 1.892 (df = 4; 155) | 1.314 (df = 12; 141) | 1.088 (df = 2; 73) | 1.376 (df = 11; 59) | 1.114 (df = 2; 81) | 0.963 (df = 11; 71) |

*Note:*
\*p<.05 \*\*p<.01 \*\*\*p<.001

## Footnotes

1 Zheng and Cohn (2011) reported the (expected) negative treatment effect on 2D:4D obtained via direct skeletal measures *in vitro*, where Huber et al. (2017) observed the opposite effect on 2D:4D measured from paw scans collected under anesthesia.

2 Ökten et al. (2002) included a small collection of hand radiographs (17 girls and 9 boys), and were unable to find differences between CAH and controls (see Ökten et al., 2002, Table 3 p. 51). However, a reanalysis of the data presented in that paper (see Richards et al., 2020, Table 4) revealed a significant difference in radiographic R2D:4D between females with CAH (n=17, M = 0.99, SD = 0.02) and female controls (n=34, M = 1.00, SD = 0.01), t = -2.393, p = 0.021, d = -0.711.

3 See https://www.cdc.gov/growthcharts/zscore.htm

4 Statistical power was calculated using G* Power 3.1 (Faul et al., 2009), for a study without repeated measures in a sample of 90 CAH and 70 Controls (as the combined sample of the current study). The availability of repeated 2D:4D measures in our sample, our reliance on measures obtained from x-rays (which are less noisy than hand scans (Ribeiro et al., 2016)) as well as our capacity to control for various confounding factors (e.g., bone age and puberty stage), was expected to increase the statistical power of our study even further. Our female sample (*N*=76) would have 90% power to detect the meta-analytic effect found in Hönekopp and Watson (2010) for the left hand of females (*d* = 0.75). Our male sample (*N*=85) would have 80% power to detect the meta-analytic effect found in Hönekopp and Watson (2010) for the left hand of males (*d* = 0.63).

5 Animal studies have shown that photographic measures of 2D:4D resulted in a higher measurement error than X-ray measurements (Lilley, et al. 2009).

